# Autopolyploid establishment under gametophytic self-incompatibility: the impact of self-fertilization and pollen limitation

**DOI:** 10.1101/2025.10.30.685356

**Authors:** Diane Douet, Xavier Vekemans, Josselin Clo

## Abstract

Polyploidy is widespread in plants, yet the establishment of neo-polyploids is limited by minority cytotype exclusion (MCE). As polyploidy has been associated with higher selfing rates in empirical studies, a shift from self-incompatibility (SI) to self-compatibility (SC) may help overcome MCE, particularly under gametophytic SI (GSI) with non-self-recognition where an automatic shift to SC is associated with polyploidy. We investigated theoretically the joint evolution of polyploidy and self-compatibility and its impact on tetraploid establishment in initially diploid populations with GSI under different pollen limitation scenarios, and with fixed or variable selfing rates for polyploids. It is shown that, for high pollen limitation scenarios, autotetraploids can invade an initially diploid population only when the selfing rates are high. Under low pollen limitation, the conditions for tetraploid invasion are less restrictive (selfing rates above 0.3). These results apply to scenarios with either fixed or variable selfing rates in polyploids, but in the latter, high mutation rates of the selfing rate are needed for polyploidy to invade. These results suggest that tetraploid establishment is possible through the evolution of selfing, without introducing a non-functional S-allele. However, the conditions strongly depend on the degree of pollen limitation.

## 1 Introduction

Polyploidy, characterized by having more than two sets of homologous chromosomes, plays a significant role in speciation and adaptation (Otto and Whitton, 2000; Van de Peer et al., 2021). It is a widespread phenomenon in plants, as about 35% of extant Angiosperms are recent polyploids (Wood et al., 2009) and many Angiosperm families show genomic signatures of ancient polyploidy events (Jiao et al., 2011; Van de Peer et al., 2017). Yet, the establishment of neo-polyploids within diploid populations is challenging for several reasons. First, polyploidization arises with several costs in neo-polyploid lineages, including genomic instability, mitotic and meiotic abnormalities, and a reduction in fitness (Comai, 2005; Otto, 2007; Doyle and Coate, 2019; Porturas et al., 2019; Clo and Kolář, 2021). In addition to these intrinsic costs of polyploidization, neo-polyploids are also facing a frequency-dependent disadvantage in mating. According to the “minority cytotype exclusion” process, reproductive opportunities for neo-tetraploids are limited due to their scarcity and the fact that fertilization between a diploid and a tetraploid results in triploid offspring that are often non-viable, sterile, or suffer from lower fertility (Levin, 1975; Ramsey and Schemske, 1998; Salony et al., 2025).

These short-term challenges raise the question of the establishment of newly formed polyploids in initially diploid populations. Many theoretical models have been proposed to address this question, offering hypotheses on how neo-polyploids could overcome minority cytotype exclusion (MCE). A first mechanism would be that new polyploid lineages are fitter than their diploid counterparts. This scenario seems unlikely under stable conditions (Porturas et al., 2019; Clo and Kolář, 2021; Salony et al., 2024), but could be reasonable during periods of environmental turmoil in which polyploids are selected in due to higher fitness in some stressful conditions (Bomblies, 2020; Van de Peer et al., 2021). Other theoretical models have focused on spatial segregation (by colonizing new habitats or by establishing at the edge of a species range for instance), as it would decrease reproductive interactions among ploidy levels, reducing the disadvantage of the minority cytotype (Baack, 2005; Griswold, 2021; Kauai et al., 2023), and favor rapid local adaptation of polyploids (Zwaenepoel, 2025). Other models have also explored the impact of different factors, such as the rate of unreduced gamete production that lead to the formation of polyploids, showing that genetic drift and environmental stochasticity can promote higher rates of unreduced gamete production and polyploids establishment (Felber, 1991; Clo et al., 2022; Gerstner et al., 2024). Finally, another class of models has investigated the evolution of different mating strategies that favor within-cytotype interactions, such as assortative mating (Husband, 2004; Oswald and Nuismer, 2011), asex-uality (Van Drunen and Friedman, 2022), or self-fertilization (Gaynor et al., 2025; Griswold, 2021; Rausch and Morgan, 2005). This article focuses on the effects of self-fertilization.

Models investigating the effect of self-fertilization found that an increased selfing rate in newly formed polyploids may facilitate their establishment, as it provides reproductive assurance (Jain, 1976), as well as a mean to reduce incoming gene flow from neighboring diploids (Antonovics, 1968). Therefore, it allows polyploid to escape MCE (Rausch and Mor-gan, 2005), even though this statement only holds when polyploids are at least as fit as their diploid counterparts (see, for example, Clo et al. (2022)), otherwise selfing polyploid lineages are eliminated in favor of fitter outcrossing diploid ones. Indeed, a relationship between self-fertilization and polyploidy has often been observed (Stebbins, 1950; Barringer, 2007). However, this relationship is still debated (Mable, 2004; Duan et al., 2024). In addition, distinctions between autopolyploids (sets of chromosomes from the same species) and allopolyploids (from different species) should be considered when interpreting these trends, as it leads to different patterns, with allopolyploid species showing higher selfing rates on average than their diploid counterparts, while autopolyploids were found to have significantly lower selfing rates than allopolyploids (Husband et al., 2008). Moreover, some studies report that polyploids experience less inbreeding depression than their diploid counterparts, at least in the early stages of establishment, potentially facilitating the evolution of selfing (Lande and Schemske, 1985; Otto and Whitton, 2000; Clo and Kolář, 2022). However, other models predict increased inbreeding depression in polyploids under certain genetic assumptions, such as how the dominance of deleterious alleles change under tetrasomic inheritance (Ronfort, 1999). These conflicting predictions highlight the complexity of the relationship between polyploidy and the evolution of mating systems.

A correlation between ploidy level and increased selfing rates could potentially involve a shift from self-incompatibility (SI) to self-compatibility (SC) in association with genome duplication (Mable, 2004). Indeed, about 45% of Angiosperm species possess a functional self-incompatibility system preventing self-fertilization in hermaphrodites (Ferrer et al., 2025).

Depending on the type of SI system, the S-phenotype of the pollen is either determined by its haploid S-genotype (gametophytic SI (GSI)) or by the diploid S-genotype of its parent plant (sporophytic SI (SSI)) (Bateman, 1952). SI systems can also be based either on self-recognition (key-lock system) or on collaborative non-self-recognition (toxin-antitoxin system) (Vekemans and Castric, 2021). Among all SI systems, one is of particular interest when considering the joint evolution of polyploidy and self-compatibility, which is the gametophytic self-incompatibility (GSI) system with collaborative non-self-recognition (Fujii et al., 2016). First, this system is the most common multiallelic SI system, already identified in 7 angiosperm families within the Eudicots (Ramanauskas et al., 2025). Second, a transition from SI to SC can be easily explained in such systems, because genome duplication mechanistically disrupts the SI reaction in heterozygous pollen through a mechanism known as competitive interaction (Lewis, 1947; Charlesworth et al., 2005; Tsukamoto et al., 2005). Indeed, tetraploid individuals mainly produce diploid pollen which, when carrying two different S-alleles, contains all the necessary anti-toxins to enable self-fertilization (Fujii et al., 2016). A more thorough explanation of the impact of whole-genome duplication on GSI systems with collaborative non-self recognition can be found in Section 1.1. However, selfing is also associated with increased homozygosity, which could potentially restore functional SI in tetraploids after some generations of self-fertilization, by producing homozygous pollen at the S-locus. The mechanisms underlying GSI are well established (Newbigin et al., 1993; Franklin-Tong and Franklin, 2003; Fujii et al., 2016), and numerous studies have focused on GSI in diploid populations, particularly regarding the conditions that maintain SI (Harkness et al., 2021; Brom et al., 2020) and the diversification of S-haplotypes (Uyenoyama et al., 2001; Gervais et al., 2014; Bod’ová et al., 2018; Stetsenko et al., 2023; Erez et al., 2024) . However, less is known about how SI systems respond to polyploidization. In particular, theoretical models are still needed to better understand how functional disruption of SI in polyploids will influence the formation and establishment of newly formed polyploids within diploid populations.

Other factors, such as pollen limitation and inbreeding depression, are considered to influence the shift from SI to SC . Indeed, studies suggest that SC is often favored under pollen-limited conditions (Charlesworth and Charlesworth, 1979; Goodwillie, 1999; Porcher and Lande, 2005). SI can become disadvantageous in such environments, because SI individuals are expected to experience an additional Allee effect, in which a reduction in population size reduces diversity at the S-locus and therefore the number of compatible mates (Wagenius et al., 2007; Leducq et al., 2010). However, this does not necessarily result in a complete tran-sition to SC (Vallejo-Maŕın and Uyenoyama, 2004). The joint evolution of self-fertilization and inbreeding depression has also been explored through theoretical models. These stud-ies show that mutations causing significant increases in selfing rates are favored by selection and can spread through populations despite high levels of inbreeding depression, thanks to the purging of deleterious or lethal mutations (Charlesworth et al., 1990; Lande et al., 1994). However, while high selfing rates provide reproductive assurance under pollen limitation, com-plete selfing is often not evolutionarily stable (Porcher and Lande, 2005). High selfing rates can lead to accumulation of deleterious mutations in small populations, potentially leading to extinction (Abu Awad and Billiard, 2017). It also can become a disadvantage due to pollen discounting, which in turn promotes the maintenance of intermediate selfing rates in populations (Holsinger, 1991; Johnston, 1998).

This work aims to better understand the consequences of the mechanistic breakdown of GSI with non-self-recognition associated with polyploidy, on the establishment of tetraploid individuals in a diploid population, under different pollen limitation scenarios. Different theoretical approaches are conducted, where the initial population is diploid and tetraploids can appear afterward through the production of unreduced gametes. The first approach is a neutral model consisting of an analytical and a simulation model. In this approach, all tetraploids are considered to be self-compatible, while diploids are strictly self-incompatible, and individuals do not have an assigned fitness, nor an S-genotype (alleles at the S-locus determining self-incompatibility). The second approach is a more complex simulation model with a GSI system with non-self-recognition, and selection according to a quantitative genetic model, in which we investigated how the mechanistic disruption of the GSI system in autotetraploid individuals, associated with changes in the selfing rate (either fixed or able to evolve through time), affects the likelihood of autopolyploid establishment.

### 1.1 GSI with collaborative non-self recognition

In gametophytic self-incompatibility (GSI), the S-phenotype of the pollen is determined by the S-genotype of its haploid pollen, whereas the two S-alleles expressed in the pistils are codominant. Collaborative non-self recognition operates as a toxin-antitoxin system. More specifically, the pistil expresses S-RNase toxins encoded by its S-alleles, which penetrate into the pollen tube and must be detoxified by antitoxins expressed by the pollen to ensure pollen tube growth and fertilization.

In diploid individuals (Figure 1(A)), pollen grains are haploid and carry all antitoxins except the one corresponding to their own S-allele. As a result, self-pollen lacks the antitoxin required to neutralize one of the pistil’s toxins and is therefore self-incompatible.

**Figure 1:**
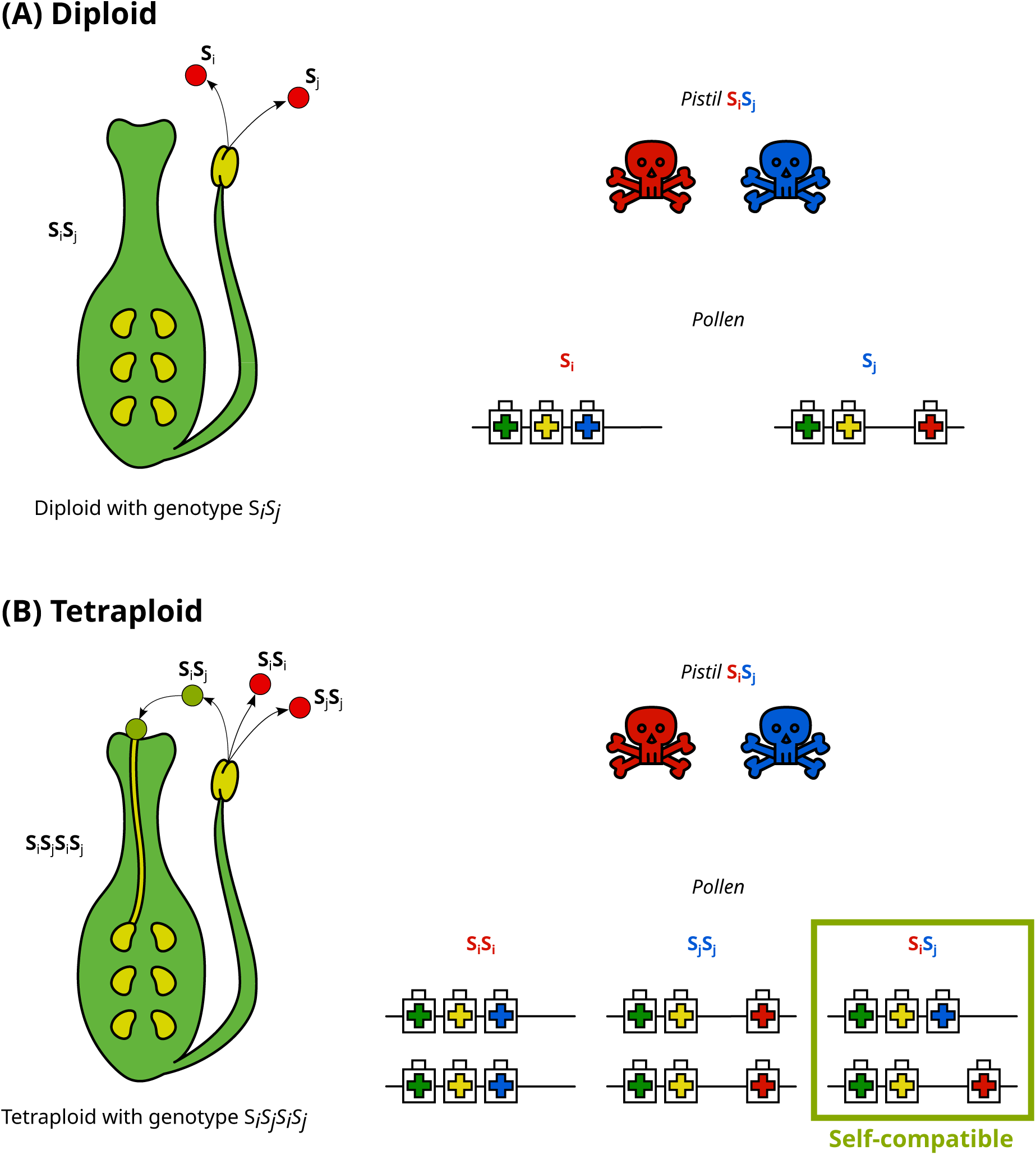
The impact of whole-genome duplication on GSI with collaborative non-self recognition. The schematics on the left represent the reproductive organs of an Angiosperm. The representation of the toxin-antitoxin system (on the right) is strongly inspired by Vekemans and Castric (2021).

**Figure 2:**
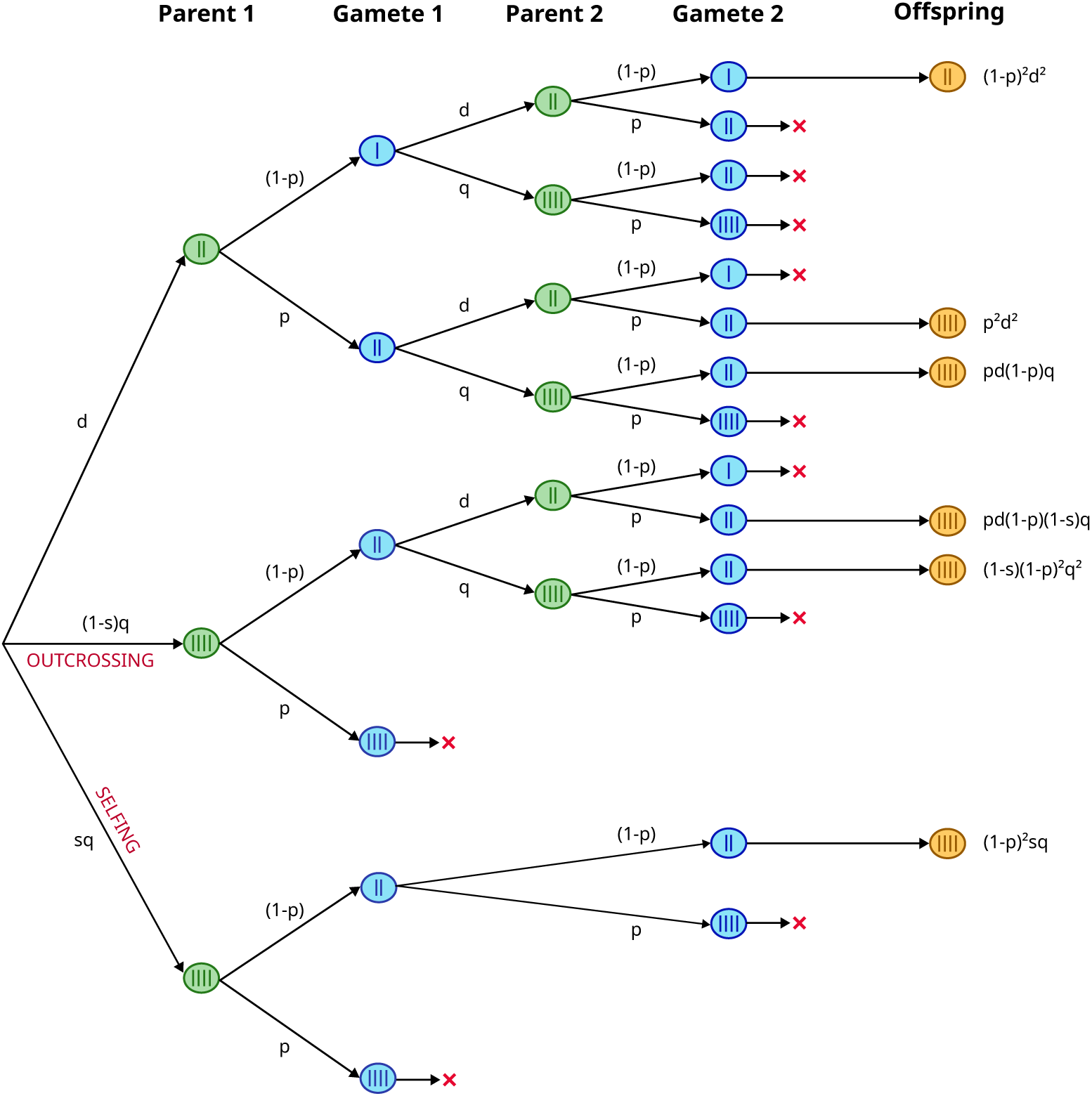
Probability tree to determine the frequencies of diploid and tetraploid individuals in the population at the next generation. A maternal parent is randomly drawn from the population and can be a diploid (obligated outcrosser), an outcrosser, or a selfing polyploid. This parent has a probability *p* of producing an unreduced gamete, and a probability (1 *p*) of producing a reduced one. If reproduction occurs through outcrossing, a pollen donor is selected from the population, either diploid or polyploid, with the same probabilities of producing reduced or unreduced gametes. In the case of selfing, the pollen comes from the maternal parent and can be either reduced or unreduced. Reproduction occurs only if both gametes are of the same size and at most diploid. Otherwise, no offspring are produced. This analytical model represents a scenario with high pollen limitation.

In tetraploid individuals (Figure 1(B)), pollen grains are diploid. If the individual is not fully homozygous at the S-locus, it produces heterozygous pollen carrying the full set of antitoxins. Such pollen is therefore self-compatible, as it can detoxify all S-RNase toxins expressed in the pistil, allowing pollen tube growth.

Consequently, in GSI with collaborative non-self recognition, there is an automatic functional shift from SI to SC associated with whole-genome duplication. Importantly, this shift does not require the introduction or fixation of a non-functional S-allele in the population, as reported in many diploid self-compatible species that have lost S-RNase expression or function (Broz et al., 2021).

## 2 Models and Methods

The notations and values used in the models and simulations are detailed in Table 1.

**Table 1:** Summary of notations and values.

| Symbol | Meaning | Default values |
| --- | --- | --- |
| $N_{\text{pop}}$ | Population size | 1000 |
| $p_u$ | Probability of producing unreduced gametes | 0.05 |
| $s$ | Selfing rate (for SC individuals) | |
| $d_n$ | Frequency of diploids in the population at generation $n$ | |
| $q_n$ | Frequency of tetraploids in the population at generation $n$ | |
| $L$ | Number of loci | 50 |
| $U$ | Mutation rate per haplotype | 0.005, 0.5 |
| $d$ | Dosage effect | 0.67 |
| $V_e$ | Variance of environmental effects | 1 |
| $\omega^2$ | Strength of selection | 1 |
| $a^2$ | Variance of mutational effects on the genome | 0.05 |
| $U_{SI}$ | Mutation rate at the S-locus | $10^{-5}$ |
| $U_{\text{SR}}$ | Mutation rate on the selfing rate | $10^{-5}, 10^{-3}, 10^{-1}$ |
| $\sigma^2$ | Variance of mutational effects on the selfing rate | 0.05, 0.1, 0.25 |

### 2.1 Neutral model

The goal is to determine the selfing rates at which tetraploids invade an initially diploid population, considering a few assumptions. The life cycle here only includes reproduction: in each generation, all adults reproduce, then die, and are replaced by the next generation of offspring. In the following models, tetraploids can appear in an initially diploid population through the production of unreduced gametes only. All diploids are considered to be self-incompatible, while all tetraploids are considered to be self-compatible. Individuals are not identified by their S-genotype, or fitness, but only by their ploidy level. In short, for fertilization to occur, the only requirement is that both gametes have the same number of chromosomes and are at most diploid.

#### 2.1.1 Analytical model

The frequency of diploids *̃d_n_*_+1_ (resp. tetraploids *̃q_n_*_+1_) in the next generation is determined by calculating the probability of drawing two haploid (resp. diploid) gametes of the population and of them meeting. The following expressions for these frequencies derive from Schematic 2:

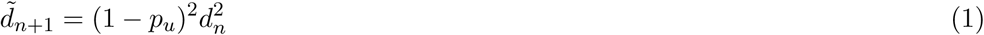

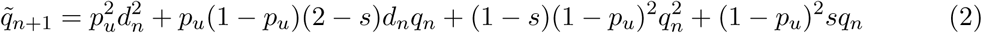

where *p_u_* is the probability of producing unreduced gametes for both diploids and tetraploids, but 4n unreduced gametes from tetraploids are lost because only diploid and tetraploid cytotypes are assumed. Finally, *s* is the “potential” selfing rate, i.e. the effective selfing rate if the individual is polyploid, while the effective selfing rate is set to zero, irrespectively of the value of *s*, if the individual is diploid. As not all gamete encounters result in a new individual for the next generation, these frequencies must be normalized by their sum to ensure that *d_n_*_+1_ + *q_n_*_+1_ = 1. Specifically, it is assumed that the number of gametes produced is always sufficient to maintain a population of size *N*_pop_ in the next generation.

#### 2.1.2 Simulation model

A simulation model is developed to observe the frequency of tetraploids at equilibrium within an initially diploid population of constant size *N*_pop_.

At the beginning of the simulation, *N*_pop_ diploid individuals are generated, each characterized only by their ploidy level. The selfing rate is fixed and is the same for each individual throughout the simulation. In each generation, the first parent is randomly selected from the population and has a probability *p_u_* of producing unreduced gametes. At this stage, the first parent can either engage in outcrossing or selfing. If the first parent is diploid, or if it is a tetraploid individual reproducing by outcrossing (which occurs with a probability 1 *− s*), a second distinct parent is randomly chosen. This second parent also has a probability *p_u_* of producing unreduced gametes. If the first parent is a tetraploid, selfing is possible as all tetraploids are considered self-compatible here. In that case, selfing occurs with a probability equal to the selfing rate *s*. A new individual is added to the next generation only if both gametes are at most diploid and have the same number of chromosomes. Otherwise, no off-spring are produced, and a new first parent is selected until the desired number of offspring *N*_pop_ is reached.

### 2.2 Gametophytic self-incompatibility (GSI) model

#### 2.2.1 General assumptions

A simulation model is built to observe the effects of self-fertilization on the maintenance of tetraploid individuals in a population of constant size *N*_pop_. This population is assumed to have a gametophytic self-incompatibility system with collaborative non-self recognition. Throughout the simulations, the self-incompatibility locus (S-locus) is assumed to contain only functional S-alleles, such that self-compatibility is only possible for tetraploid individuals that are not fully homozygous at the S-locus (cf Section 1.1). The population is initially diploid, and tetraploid individuals can appear in the population through the production of unreduced gametes. Higher levels of ploidy (such as hexaploids) are not considered here. Each individual has several S-alleles and several chromosomes equal to its ploidy level. Each chromosome contains *L* loci used to compute individual fitness, each locus possibly containing an infinite number of alleles. Each locus stores a real value, initially set to zero, which can mutate to another independent and random value (allele) during the simulation. The selfing rate is either fixed and the same for all individuals, or it can evolve during the simulation. In the case where the selfing rate is allowed to evolved, the selfing rate is represented as one single value in each individual. From a modeling perspective, the selfing rate is equivalent to an individual trait encoded by an infinite number of loci, whose value is defined as the average effect of these loci. The selfing rate is a “potential selfing rate”, i.e. it represents the effective selfing rate if the individual is self-compatible, while the effective selfing rate is set to zero, irrespectively of the value of *s*, if the individual is self-incompatible. Moreover, the model assumes prior selfing (Lloyd, 1979), meaning that autonomous selfing takes place just before anthesis so that the effective selfing rate does not depend on the availability of compatible pollen.

The fitness of an individual is computed using a quantitative genetic model similar to the one in Clo (2022). The phenotypic trait value *z* of an individual is defined as:

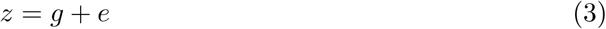

where *e* is a random environmental effect drawn from a normal distribution with mean 0 and variance *V_e_*, and *g* denotes the genotype of an individual, calculated as follows:

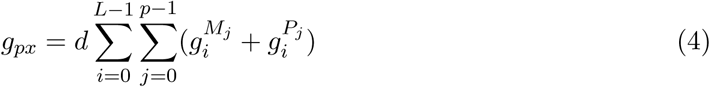

where 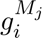 (resp. 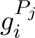 ) is the value stored in the locus *i* of the maternal (resp. paternal) chromosome *j*, *p* is the ploidy level, and *d* is the dosage factor accounting for the effect of ploidy on genotypic values (set to 1 for diploids). In other words, the genotype of an individual is the sum of all values stored in its genome, scaled by the dosage factor for polyploid populations.

The fitness *W*_(*z*)_ of an individual with phenotype *z* is given by:

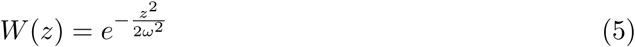

where *ω*^2^ denotes the width of the fitness function, i.e. the strength of selection.

#### 2.2.2 Simulation model

At the beginning of the simulation, *N*_pop_ identical diploid genomes are created with 2 chromosomes per individual. The value initially stored in each locus is the optimum value 0. S-genotypes are also created, with 2 functional S-alleles per individual randomly chosen among all possible 100 initial S-alleles. A “potential” selfing rate is also attributed to each individual (as one single value, equivalent to an individual trait encoded by an infinite number of loci), which is the same for all in the case of a fixed selfing rate *s*, and 0 in the case where the selfing rate can evolve. This selfing rate is only realized for self-compatible (SC) tetraploid individuals, whereas self-incompatible (SI) individuals are obligate outcrossers independently of the value of the parameter.

The next generation is obtained as follows: a first maternal parent is selected with a probability relative to its fitness *W* (*z*), and produces unreduced gametes with probability *p_u_*, as all individuals in the population. If this parent is diploid (and hence SI), it necessarily engages in outcrossing. In such a case, a distinct paternal parent is selected (with a probability relative to its fitness), as well as its pollen grain. The pollen’s S-allele(s) is then compared with the two S-alleles of the mother plant. If the pollen and pistil S-alleles are compatible, a new individual is added to the next generation as long as the gametes have the same number of chromosomes and are at most diploid. Otherwise, no new individual is added to the population, and the process is repeated with a new paternal parent until fertilization is achieved or until the set number of attempts with the same maternal plant is exceeded (1, 5, or 20 attempts). If the maternal parent is tetraploid, it can either engage in outcrossing following the same procedure as for diploids, or engage in self-fertilization with a probability equal to its selfing rate. In such a case, the S-alleles of the maternal pistil and of the self pollen are compared. If the S-alleles are compatible, and if the gametes have the same number of chromosomes and are at most diploids, a new selfed individual is added to the next generation. Otherwise, no new selfed individual is added to the population, and the process is repeated with a new self pollen grain until fertilization is achieved or until the set number of attempts with the same maternal plant is exceeded (1, 5, or 20 attempts). The number of attempts mimics pollen limitation scenarios (Vekemans et al., 1998), 1 attempt being the maximal limitation (i.e. one ovule receives a single pollen grain), and 20 attempts modeling low pollen limitation (i.e. one ovule receives up to 20 pollen grains before being rejected). If the number of attempts is exceeded, a new maternal parent is selected, and the process continues until the desired population size *N*_pop_ is reached. Each parent passes on a gamete to the next generation, and one (for diploids) or two (for tetraploids) of their alleles at all sites (the *L* fitness loci and the S-locus), depending on the size of the gamete. Crossovers are modeled here as a random permutation between alleles present at the same locus on homologous chromosomes, i.e. free recombination among adjacent loci. The selfing rate of the new individual is the average of the selfing rates of its parents, as expected under the hypothesis that selfing rate is a quantitative trait. Next, mutations are introduced within the genome of the new individual formed.

A simulation run consists of three phases in which different types of mutation occur. In the first phase (2 000 generations), all individuals remain diploids and are subject to mutations in their S-genotypes. To be more precise, each S-allele can mutate to one the *k −* 1 (*k* = 100) other possible S-alleles with probability *U*_SI_. The number of mutations is drawn from a binomial distribution with parameters (*N*_pop_ *×* ploidy*, U_SI_*), and their positions are chosen at random. In the case where the selfing rate can evolve, mutations on an individual’s selfing rate are also introduced in the first phase. The number of mutations is drawn from a binomial distribution with parameters (*N*_pop_*, U_SR_*), and the individuals subject to the mutation are randomly chosen among the population. The additive value of the mutation is drawn from a normal distribution with parameters 0 and *σ*^2^, controlling the speed of evolution toward higher selfing rates. Selfing rates are restricted to remain between 0 and 1, which can create boundary effects. However, we expect this bias to have a limited impact on the results, causing mainly a slight shift away from values close to 0 or 1. In the second phase (2 000 generations), mutations to the *L* fitness loci coding for the quantitative traits under selection can also occur. The number of mutations per haplotype is drawn from a Poisson distribution with parameter *U* , and their positions in the genome are randomly sampled from the *L* possible loci. The additive value of this mutation is then drawn from a normal distribution with parameters 0 and *a*^2^. In the last phase (200 000 generations), unreduced gametes are introduced during the reproduction phase with probability *p_u_*, and tetraploid individuals can appear in the population.

At the end of the simulation, the frequency of tetraploids in the population is computed, as well as three other frequencies: the frequency of SC individuals, the frequency of SC pollen (i.e. heterozygous pollen) produced by these individuals, and the frequency of individuals engaging in selfing averaged over the last 10 000 generations. Inbreeding depression is also computed using the following formula:

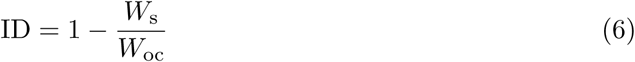

where *W*_s_ is the fitness of selfed individuals, and *W*_oc_ the fitness of randomly outcrossed individuals in the population. Each fitness is computed by sampling 100 individuals in the population. The fitness values of diploids (after reaching an equilibrium, before introducing unreduced gametes in the population) and of tetraploids (at the end of the simulation) are also outputs of the simulation model.

## 3 Results

### 3.1 Neutral model

#### 3.1.1 Analytical results Equilibria

Using the fact that *d_n_*_+1_ = *d_n_*at equilibrium and *q_n_* = 1*−d_n_* for all *n*, we find two equilibria *{d*_eq1_*, q*_eq1_*}* = *{*0, 1*}* and *{d*_eq2_*, q*_eq2_*}* = *{d , q }* with:

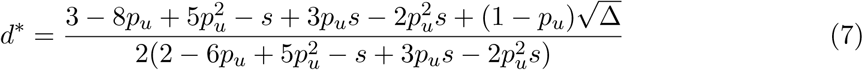

where 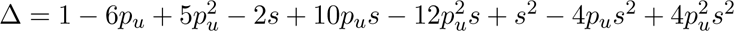 is the determinant, and *q*^∗^ = 1 *− d*^∗^.

As *d*^∗^ is obtained as the solution of a second-order polynomial equation, the second equilibrium *{d*^∗^*, q*^∗^*}* is valid as long as Δ *≥* 0. If Δ *<* 0, the only possible equilibrium is *{*0, 1*}*, which implies that tetraploids have established into the population. Therefore, the threshold selfing rate at which tetraploids establish into the initially diploid population is the solution to the equation Δ = 0, which is the following:

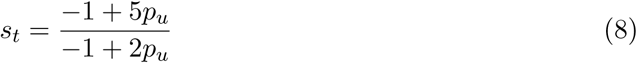

Using the equations above, we obtain the frequency of tetraploids in the population at equilibrium as a function of the selfing rate:

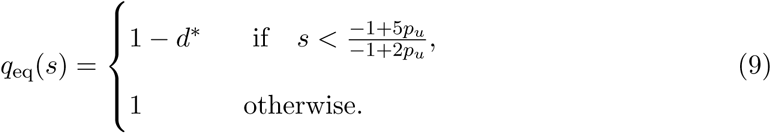

To validate the analytical expressions for the threshold selfing rate and the number of tetraploids at equilibrium, we conducted both numerical and analytical analyses for comparison. Figure S1 confirms the accuracy of these expressions.

##### Stability

In order to determine the stability of these equilibria, we use a standard stability analysis (see e.g. (Otto and Day, 2011, Chapter 5)) and *Wolfram Mathematica*. First, we differentiate the function *f* (*d_n_*) = *d_n_*_+1_. Since we consider discrete generations here, an equilibrium *d*_eq_ is stable if *|f* ^′^(*d*_eq_)*| <* 1.

Because *f* ^′^(0) = 0 for all values of *p_u_*and *s*, the first equilibrium *{d*_eq1_*, q*_eq1_*}* = *{*0, 1*}* is always stable, regardless of the parameter values.

For the second equilibrium, we found that *|f* ^′^(*d*^∗^)*| <* 1 holds when

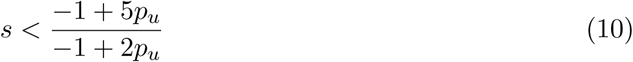

and when the rate of unreduced gamete production *p_u_* remains below 0.2, a range considered realistic for this type of biological system. Note here, that the equilibrium *{d*_eq2_*, q*_eq2_*}* = *{d , q }* is stable when *s < s_t_*, which is precisely its range of validity. In other words, when *p_u_ <* 0.2, the equilibrium *{d*^∗^*, q*^∗^*}* is always stable when it exists.

The equilibrium reached by the system depends on the value *s* encoded at the selfing rate locus and on the initial conditions. Specifically, the equilibrium (*d*^∗^*, q*^∗^) does not hold when *s ≥ s_t_*, whereas the equilibrium (0, 1) (corresponding to tetraploid establishment) is always stable. In contrast, since we consider initially diploid populations here, i.e. *d*_0_ = 1 and *q*_0_ = 0, the system converges toward the equilibrium (*d*^∗^*, q*^∗^) for selfing rates below the threshold *s_t_*. This suggests that the probability of tetraploid establishment is zero when selfing rates are below this threshold.

#### 3.1.2 Simulation results

Figure 3 presents the frequency of tetraploids in the population at equilibrium using the neutral model. First, the results obtained with the analytical model match those obtained using the individual-based simulations for selfing rates lower than 0.8 (Figure 3). Therefore, the frequency of tetraploids at equilibrium depends exclusively on the probability of producing unreduced gametes *p_u_* according to Section 2.1.1. However, the threshold selfing rate for which tetraploids invade the population obtained with the simulation model is slightly lower than the one obtained analytically (Figure 3).

**Figure 3:**
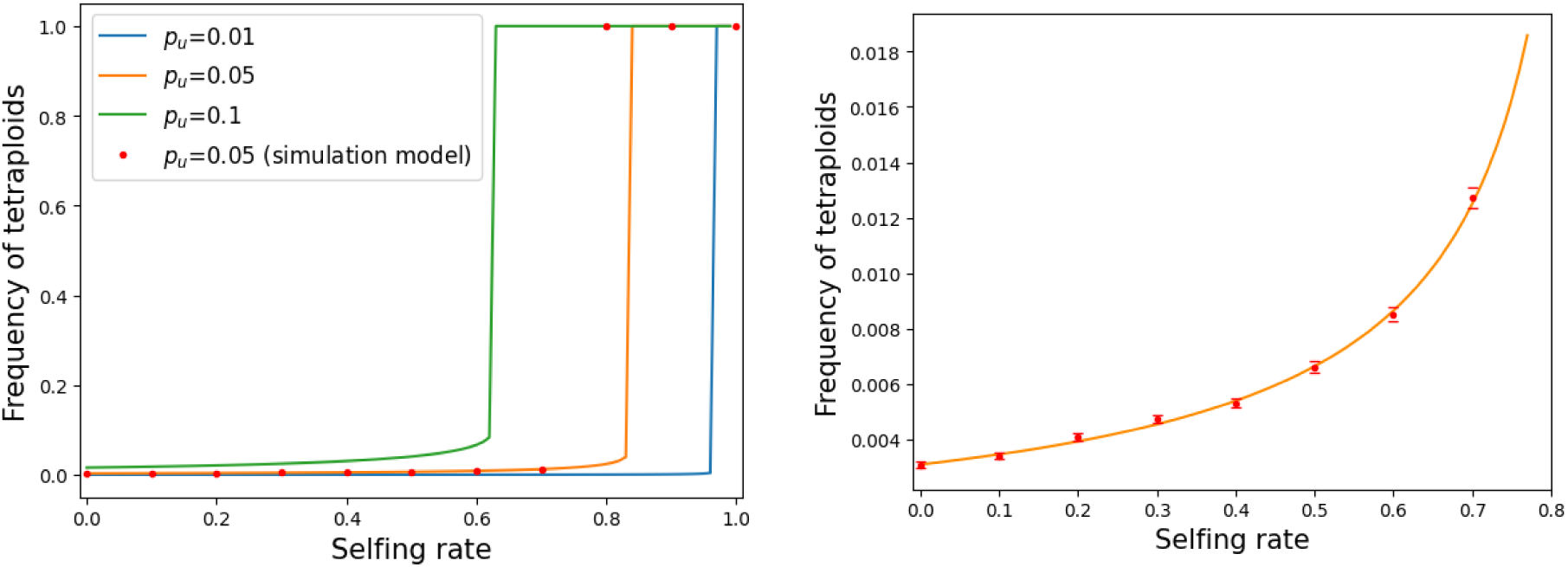
Frequency of tetraploids at equilibrium as a function of the selfing rate for different values of the probability of producing unreduced gametes pu obtained using the neutral model (left), and a zoomed version for *p_u_* = 0.05 (right). The full lines represent the results obtained with the analytical model, while the red dots are obtained using the simulation model (50 runs) with *N_pop_* = 1000, and the red dashes represent the 95% confidence intervals.

Moreover, it can be observed that the higher the probability of producing unreduced gametes, the lower the selfing rate threshold allowing tetraploids to invade, which is expected. More importantly, Figure 3 shows that tetraploids are able to establish in the initially diploid population, only when the selfing rate is quite high. Focusing on *p_u_* = 0.05, the selfing rate needs to be higher than 0.8 for the tetraploids to invade the population. Tetraploids will invade the population when the selfing rate is high enough for them to overcome minority cytotype exclusion, a threshold selfing rate that depends only on the rate of unreduced gametes.

### 3.2 GSI model

#### 3.2.1 Fixed selfing rates

We explored different scenarios of pollen limitation, where the first maternal parent is rejected after 1, 5 or 20 unsuccessful reproduction attempts. Under high pollen limitation, tetraploid invasion occurs only when selfing rates exceed 0.8 (Figures 4a and S2a), which is consistent with the results from the neutral model (Figure 3). In contrast, under low pollen limitation, tetraploid invasion occurs at much lower selfing rates (Figure 5a). Specifically, when selfing rates exceed 0.3, tetraploids always invade the population in our simulations. Even at a selfing rate of 0.1, tetraploid invasion was observed in a few simulations, and in 30% of the simulations at a selfing rate of 0.2. The most striking difference between the high pollen limitation scenarios (Figures 4 and S2) and the low pollen limitation scenario (Figure 5) lies in the value of the threshold selfing rate required for tetraploids to invade an initially diploid population. This difference between pollen limitation scenarios can be explained as follows: the higher the number of attempts before rejecting the maternal parent, the higher the likelihood of finding a compatible mate under challenging conditions, e.g. finding a diploid pollen grain (either an unreduced gamete from a diploid or a reduced gamete from a tetraploid individual) to fertilize a diploid ovule produced by a tetraploid individual.

**Figure 4:**
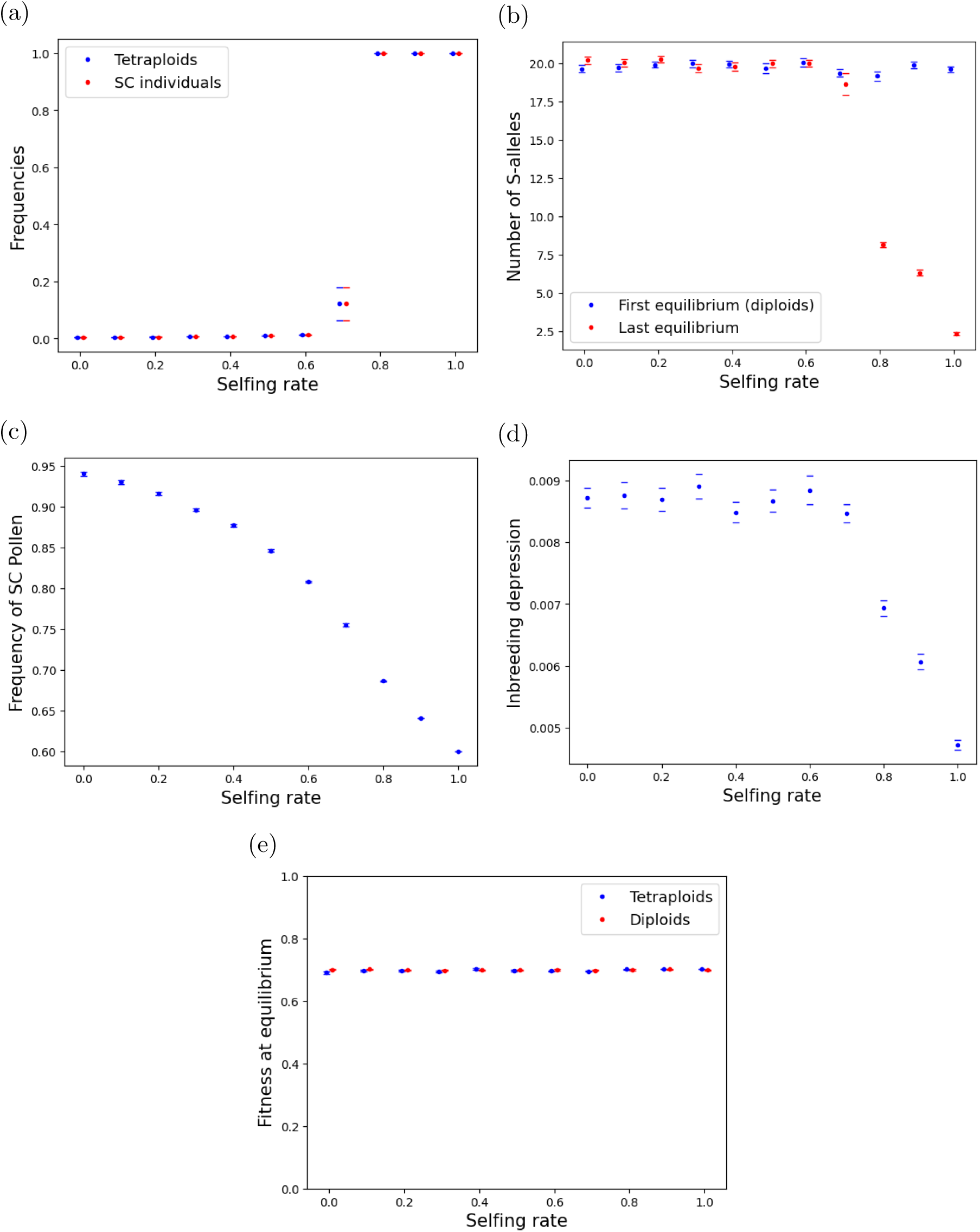
Evolutionary dynamics under high pollen limitation. (a) Frequencies of tetraploids and SC individuals, (b) number of S-alleles at equilibrium for diploids and at the end of the simulation, (c) frequency of SC pollen produced by SC individuals, (d) inbreeding depression over the last 10000 generations, and (e) fitness of diploids and tetraploids at equilibrium when the selfing rate is fixed. Error bars stand for 95% confidence intervals. Results of 30 simulation runs under high pollen limitation (1 attempt) for each selfing rate using the parameters detailed in Table 1, specifically *U* = 0.005, *ω*^2^ = 1, *N*_pop_ = 1000 and *p_u_* = 0.05

**Figure 5:**
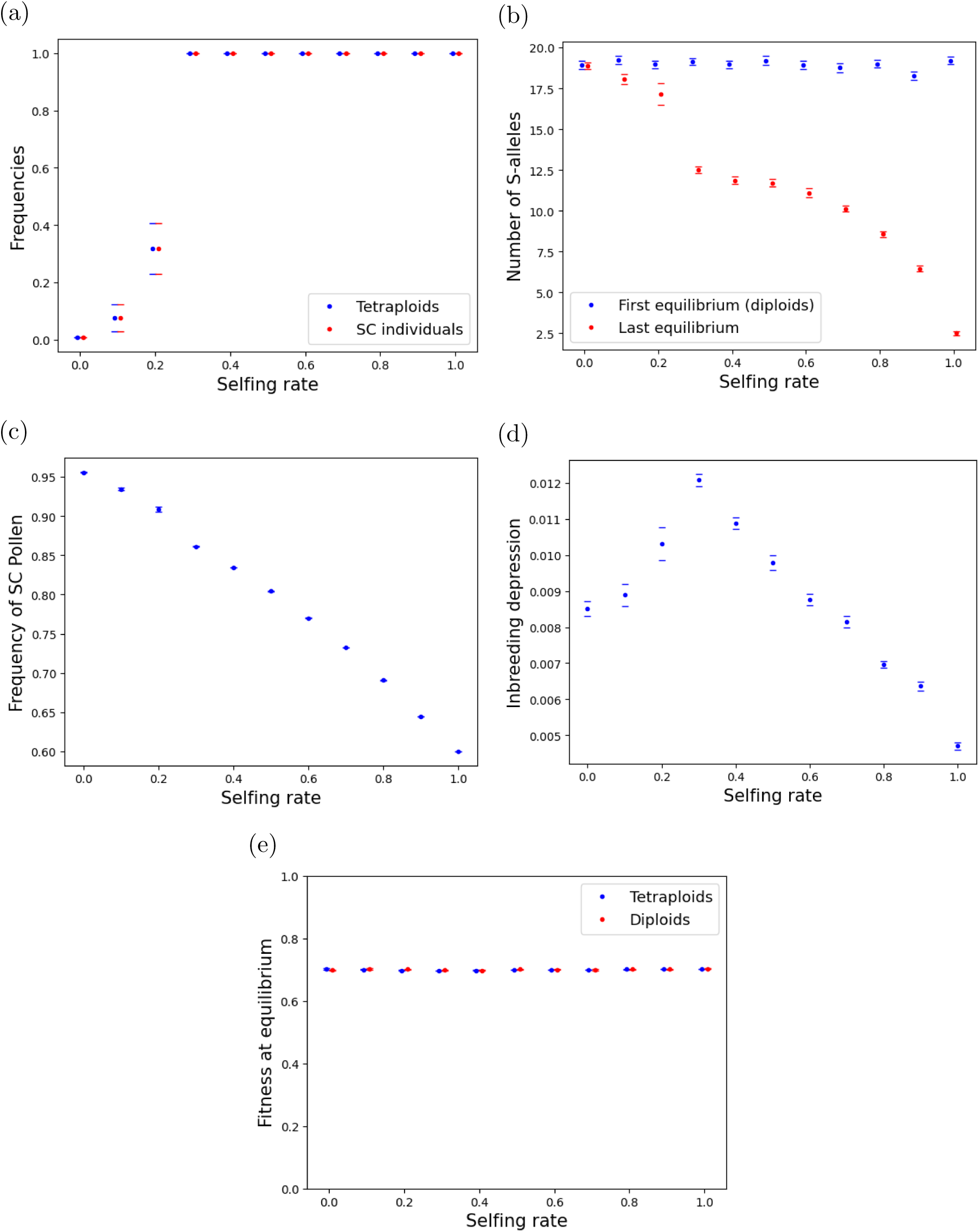
Evolutionary dynamics under low pollen limitation. (a) Frequencies of tetraploids and SC individuals, (b) number of S-alleles at equilibrium for diploids and at the end of the simulation, (c) frequency of SC pollen produced by SC individuals, (d) inbreeding depression over the last 10000 generations, and (e) fitness of diploids and tetraploids at equilibrium when the selfing rate is fixed. Error bars stand for 95% confidence intervals. Results of 30 simulation runs under low pollen limitation (20 attempts) for each selfing rate using the parameters detailed in Table 1, specifically *U* = 0.005, *ω*^2^ = 1, *N*_pop_ = 1000 and *p_u_* = 0.05.

Nonetheless, the different pollen limitation scenarios exhibit similar patterns. Even though all tetraploids are SC, the frequency of SC pollen produced by SC individuals decreases with increasing selfing rate (Figures 4c, S2c, and 5c), revealing a drop in heterozygosity at the S-locus. Furthermore, in cases where tetraploids invade the population, an increase in the selfing rate is also associated with a decrease in both the number of S-alleles maintained within the population (Figures 4b, S2b, and 5b) and the level of inbreeding depression at the end of the simulation (Figures 4d, S2d, and 5d). These last observations, i.e. a decrease in heterozygosity, a loss of diversity, and a lower inbreeding depression, remain unchanged under different pollen limitation conditions and are consistent with what is expected for highly selfing populations. In addition, Figures 4e, S2e, and 5e show no difference between the fitness of diploids and tetraploids at equilibrium, which explains the consistency of the results under high pollen limitation with those from the neutral model.

##### Higher levels of inbreeding depression

Additional simulations were performed using a high mutation rate per haploid genome (*U* =0.5). The resulting values of inbreeding depression are consistent with empirical observations, which indicate that polyploids exhibit inbreeding depression of 0.1 in neo-polyploids and around 0.3 in natural polyploids on average (Clo and Kolář, 2022). Under low pollen limitation, the results are quantitatively the same as for lower mutation rate (Figure S4). Figure S3 shows that under high pollen limitation, tetraploid establishment occurs at lower selfing rates when the mutation rate is high (*U* = 0.5) than when it is low (*U* = 0.005). This can be explained by the fact that tetraploids exhibit higher fitness than their diploid progenitors (Figure S3e) as they more effectively mask deleterious mutations. Therefore, they have an increased probability of persisting and contributing to the next generations, whereas they are more likely to be lost through genetic drift at lower mutation rates. When inbreeding depression is high, self-fertilization represents an opportunity for tetraploids to maintain even at lower selfing rates. For this reason, we chose to emphasize the low inbreeding depression scenario in this article, as it represents a more conservative framework where tetraploid establishment is more challenging.

#### 3.2.2 Evolving selfing rates

When the selfing rate (or self-pollen rate for diploids) of each individual is allowed to evolve, Figures 6, S5, and 7 show that tetraploid invasion depends mainly on the mutation rate *U*_SR_ and on the extent of pollen limitation. For a realistic mutation rate of *U*_SR_ = 10^−5^, tetraploids never invade the population, regardless of the pollen limitation scenario and of the variance of mutational effects on the selfing rate *σ*^2^ (Figures 6a, S5a, and 7a). Although some individuals with high selfing rates (above 0.8) appear in the population, their frequency remains too low to enable tetraploid invasion. At higher mutation rates (*U*_SR_ = 10^−3^ and *U*_SR_ = 10^−1^), individuals reach a wide range of selfing (or self-pollen) rates between 0 and 1. However, under high pollen limitation, tetraploids never invade the population, regardless of the other parameters (Figures 6b, 6c, S5b, and S5c). In contrast, Figures 7b and 7c indicate that tetraploid invasion always occurs under low pollen limitation when the mutation rate *U*_SR_ is not too low, regardless of the variance *σ*^2^, suggesting once again that the absence of pollen limitation plays a key role in tetraploid establishment.

**Figure 6:**
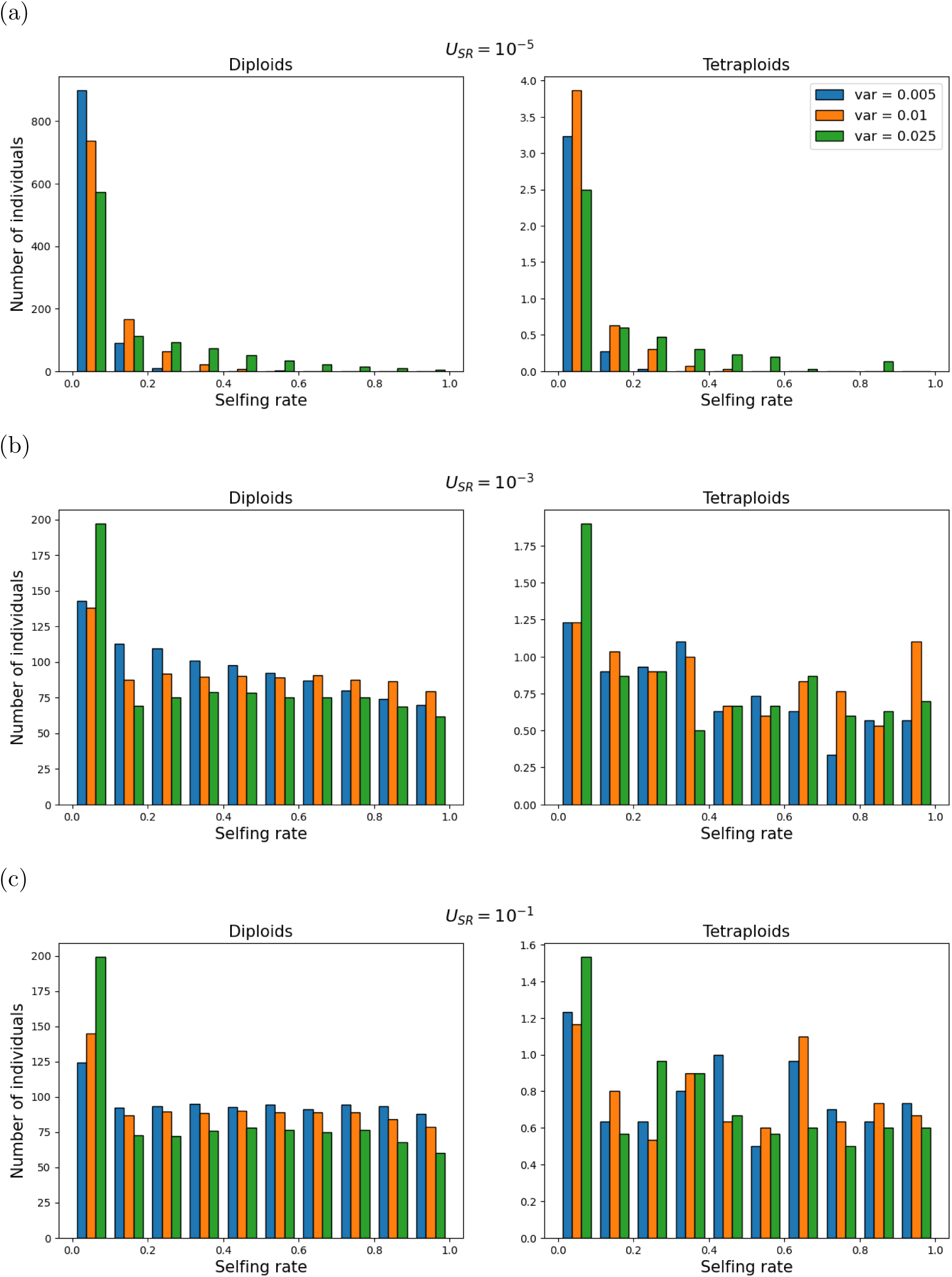
Distribution of the selfing rates of diploids (left) and tetraploids (right) at the end of the simulation for three different values of the variance *σ*^2^ = 0.05, 0.1 and 0.25 and for different values of the mutation rate: (a) *U*_SR_ = 10^−5^, (b) *U*_SR_ = 10^−3^ and (c) *U*_SR_ = 10^−1^. Results averaged over 30 simulation runs under high pollen limitation (1 attempt) when the selfing rate can evolve using the parameters detailed in Table 1, specifically *N*_pop_ = 1000 and *p_u_* = 0.05.

**Figure 7:**
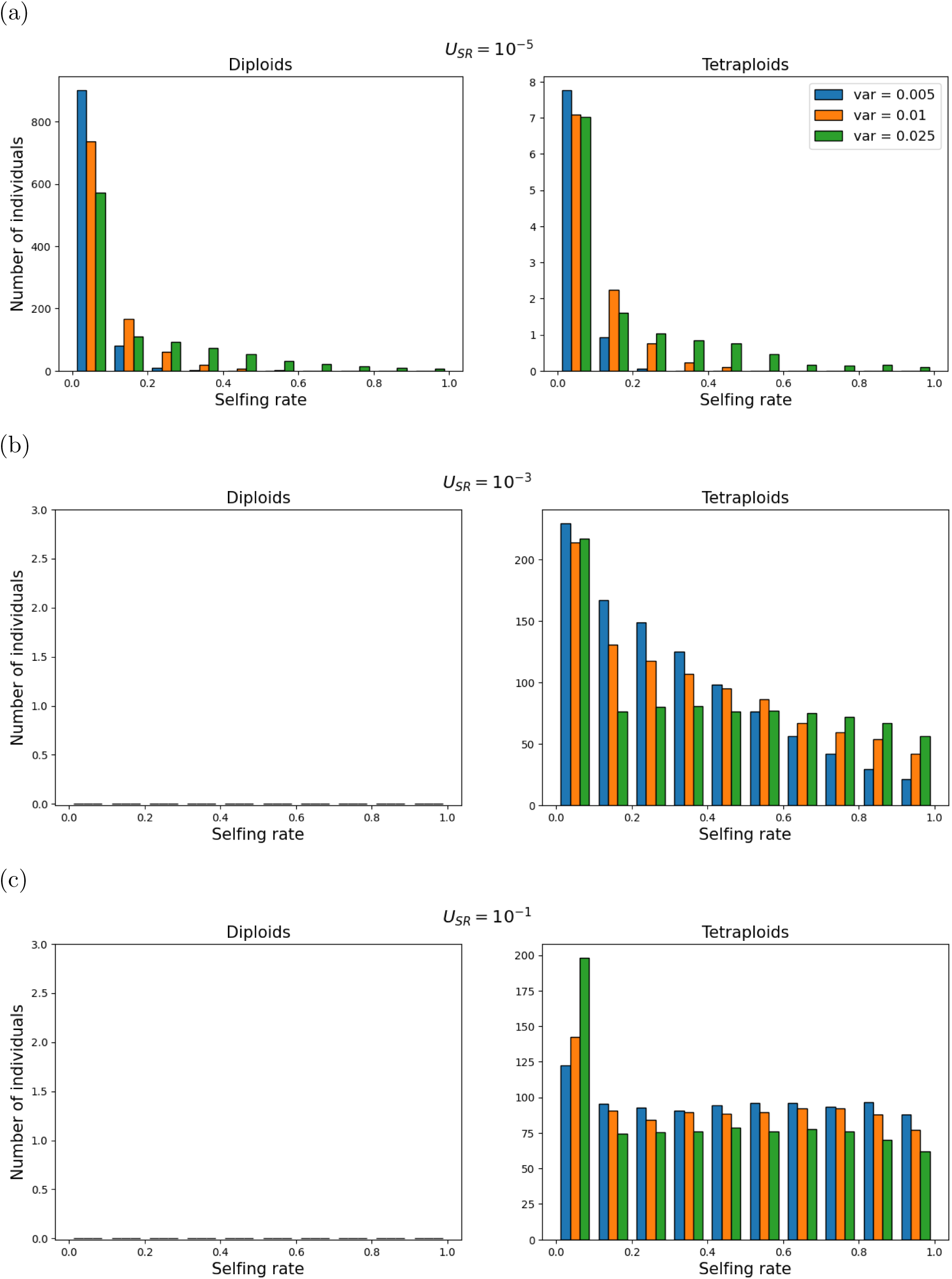
Distribution of the selfing rates of diploids (left) and tetraploids (right) at the end of the simulation for three different values of the variance *σ*^2^ = 0.05, 0.1 and 0.25 and for different values of the mutation rate: (a) *U*_SR_ = 10^−5^, (b) *U*_SR_ = 10^−3^ and (c) *U*_SR_ = 10^−1^. Results averaged over 30 simulation runs under low pollen limitation (20 attempts) when the selfing rate can evolve using the parameters detailed in Table 1, specifically *N*_pop_ = 1000 and *p_u_* = 0.05.

## 4 Discussion

In this paper, we investigated the impact of self-fertilization on the establishment of tetraploids within initially diploid populations under a GSI system with non-self-recognition. A key functional property of such SI systems is that whole-genome duplication results in an automatic shift from SI to SC, without the need to introduce a non-functional S-allele (Fujii et al., 2016). This has been confirmed by empirical studies in *Solanaceae* (Robertson et al., 2011), exhibit-ing a GSI system functioning as a collaborative non-self recognition system producing a pistil S-RNase toxin. A classical exception to this rule has been suggested in the literature for the *Prunus* genus in *Rosaceae*, where polyploid *Prunus* have been shown to maintain a functional GSI system (Hauck et al., 2006; Robertson et al., 2011), but it has now been demonstrated that the *Prunus* GSI system, despite relying on a S-RNase female component, is indeed function-ing as a self-recognition system (Aguiar et al., 2015; Wang et al., 2025). Our results indicate that tetraploid establishment is possible through the evolution of self-fertilization alone, i.e. without introducing an extrinsic fitness advantage of polyploidy, although the conditions for invasion vary depending on the degree of pollen limitation and the rate of unreduced gamete production.

### Evolution of selfing and tetraploid establishment

Under high pollen limitation, we found that tetraploid invasion is restricted to cases where the selfing rate of the entire population is high (above 0.8 for biologically realistic values of the rate of unreduced gamete production, i.e. *p_u_ ≤* 0.05), and does not occur through the evolution of selfing, regardless of parameter values. Although the selfing rate threshold required for tetraploid invasion is high, it is comparable to values observed in many polyploid species. For instance, Barringer (2007) reported that polyploid annuals have a mean level of self-fertilization close to 0.8, while herbaceous and woody perennials exhibit lower rates on average (around 0.5 and 0.3, respectively). Our threshold selfing rate is also in line with the average outcrossing rate of 0.2 found in allopolyploids by Husband et al. (2008), while autopolyploids show significantly higher outcrossing rates on average (*t* = 0.64) though exceptions exist - for example, *Townsendia hookeri* autopolyploids have an outcrossing rate as low as 0.03 (even if the low outcrossing rate can also be explained by the apomictic reproductive system of the species (Thompson and Whitton, 2006)). Although our model mimics a situation corresponding to autopolyploidy, the low selfing rates reported by Husband et al. (2008) for autopolyploids may not be relevant here because they did not sample species with a non-self recognition GSI system that promotes functional loss of SI. Regarding allopolyploids, a similar functional shift from SI to SC could be expected under GSI with collaborative non-self recognition (Duan et al., 2024). Indeed , a high degree of allele sharing between parental species is expected in multiallelic SI systems (Vekemans and Slatkin, 1994), poten-tially allowing pollen antitoxins to function across parental species subgenomes. However, as suggested by Harkness and Brandvain (2021), if the two parental species are too divergent, certain antitoxins might have been lost in one of the species, preventing detoxification of all S-RNase toxins in the other species, therefore leading to self-incompatible allopolyploids. As discussed by Novikova et al. (2023), some allopolyploids can be formed by hybridization between an outcrossing species with functional SI and a selfer lacking functional SI. In such a case, the non-functional S-allele present in one subgenome can produce an SC phenotype for the neo-allopolyploid individuals. Further empirical data on the mating system of polyploid taxa discriminating among auto- and allopolyploidy are crucially needed.

Under low pollen limitation, our results indicate that tetraploid invasion occurs from much lower selfing rates (above 0.3, and even for selfing rates as low as 0.1 and 0.2 in some simulation runs). When selfing rates are allowed to evolve, tetraploids successfully invade the population under low pollen limitation as long as the mutation rate *U*_SR_ is sufficiently high (*≥* 10^−3^). Since selfing provides reproductive assurance, one might expect SC tetraploids to be more favored in pollen-limited environments. However, our results suggest the opposite: a realistic evolution of the selfing rate following whole-genome duplication (i.e. selfing rates going from 0 to 0.3 *−* 0.5) would be more likely in stable conditions with reasonably large populations (low pollen limitation scenario) than in colonization conditions with strong bottlenecks and fewer potential mates (high pollen limitation scenario). This seems to contradict previous studies predicting that pollen-limited environments favor the evolution of self-fertilization toward high selfing rates (Porcher and Lande, 2005; Vallejo-Maŕın and Uyenoyama, 2004). However, these studies focus exclusively on diploid populations, while those that address tetraploid establishment typically examine other factors, such as environmental constraints or pollen dispersal (Griswold, 2021), rather than scenarios where pollen is limited. Our results can be explained as follows: under low pollen limitation, tetraploids can also rely on outcrossing to overcome MCE as they are more likely to receive compatible diploid pollen, either from diploid donors producing unreduced gametes or from other tetraploid individuals. In contrast, under strong pollen limitation, they need to rely almost exclusively on self-pollen to receive diploid pollen, which requires higher values of the selfing rate. Overall, these findings align with theoretical models suggesting that higher selfing rates increase the likelihood of polyploid persistence (Baack, 2005; Rausch and Morgan, 2005). However, Clo et al. (2022) showed that self-fertilization can negatively impact tetraploid establishment when polyploidy is associated with a fitness cost. Note that diploids and tetraploids exhibit no differences in fitness in our simulations.

### S-allele diversity, homozygosity and inbreeding depression

We also found that the diversity of S-alleles in the population is impacted by tetraploid establishment. In diploids, the number of S-alleles is maintained by strong negative frequency-dependent selection, as a pollen with a rare S-allele is more likely to find a compatible mate compared to a pollen with a frequent S-allele (Wright, 1939). Our results for the number of S-alleles maintained at equilibrium in the diploid population are consistent with the estimate given by the formula in Yokoyama and Nei (1979), which predicts approximately 20 alleles for a population of our size under GSI. When tetraploid SC individuals invade the population, the number of S-alleles present in the population decreases with increased selfing rates, reflecting a loss of diversity at the S-locus. This pattern is consistent with the fact that selfing is associated with a reduction in genetic diversity in general, especially in highly selfing populations (Charlesworth and Charlesworth, 1995; Lande and Porcher, 2015; Abu Awad and Roze, 2018), and with the fact that the negative frequency-dependent selection acting on the S-locus is relaxed due to functional disruption of the SI reaction under polyploidy.

However, a minimal number of S-alleles still needs to be maintained to ensure SC (two S-alleles in that case), since fully homozygous tetraploids at the S-locus would revert to SI and could not self-fertilize anymore. In our simulations, selfing reaches very high levels when considering functional S-alleles only, i.e. without introducing a non-functional S-allele into the population. That is specific to GSI with non-self-recognition, as polyploidization automatically renders the SI system leaky. This contrasts with sporophytic SI systems, such as those in the Brassicaceae, where the breakdown of SI requires the presence of a non-functional S-allele (Vekemans et al., 2014; Novikova et al., 2023). From an empirical point of view, the recent sequencing of a pistil transcriptome of a polyploid *Rebutia pygmaea* (Cactaceae) revealed the presence of at least four S-RNase alleles (B. Igic, personal communication). In addition, more than two S-alleles have also been reported in some tetraploid individuals of *Lycium parishii* (Solanaceae) (Savage and Miller, 2006). However, the self-incompatibility status of polyploid individuals and the functionality of their S-alleles have not been established in these two cases. Hence, due to limited empirical evidence from polyploid species with a collaborative non-self recognition GSI system, it remains unclear whether such species can maintain functional S-alleles following polyploidization, or if they eventually fix a non-functional S-allele after an intermediate phase. Further research is needed on the subject to fill this knowledge gap, for instance through large-scale sequencing of S-RNases from recent polyploids. More generally, studying the difference in average selfing rates in auto- and allopolyploid populations or species compared to their diploid progenitors in plant families with a non-self recognition GSI system would be highly informative.

On a different note, higher selfing rates are also associated with greater homozygosity (Burgarella and Gĺemin, 2017), which is consistent with our observation of reduced heterozygosity at the S-locus among SC individuals as selfing increases. Our simulations also show that inbreeding depression declines with rising selfing rates, which is expected as increased ho-mozygosity facilitates purging of deleterious alleles (Ronfort, 1999; Charlesworth and Willis, 2009).

### Perspectives

Our study focuses on selfing as a strategy to overcome MCE, but other factors can also promote the establishment and persistence of polyploids. For example, we did not model reproductive isolation in outcrossing, such as assortative mating, which could enhance the chances of tetraploid establishment (Oswald and Nuismer, 2011). Polyploidy is often associated with differences in floral traits and flowering time, which can lead to reproductive isolation, either by attracting different pollinators or by reducing overlap in reproductive periods (Segraves and Thompson, 1999; Vamosi et al., 2007). More generally, strong assortative mating hinders the exclusion of tetraploids in populations where unreduced gametes are in-troduced, therefore facilitating their ability to overcome MCE (Husband and Sabara, 2004). We also assumed that triploid individuals are sterile or non-viable, a common assumption in theoretical models (Levin, 1975; Ramsey and Schemske, 1998; Felber, 1991), and supported by empirical studies reporting reduced fitness in triploid individuals (Husband and Sabara, 2004; Burton and Husband, 2000). However, both theoretical and empirical work suggest that the presence of a triploid bridge can facilitate tetraploid establishment, depending on the fitness of triploid individuals (Husband, 2004; Köhler et al., 2010; Schinkel et al., 2017; Felber and Bever, 1997). Apart from potential extensions, the model could also be adapted to other types of self-incompatibility systems, such as GSI with self-recognition or sporo-phytic self-incompatibility (SSI), which do not lead to automatic functional breakdown under polyploidy (Hauck et al., 2006; Duan et al., 2024). In those systems, the breakdown of SI necessarily occurs through the spread of a mutated non-functional SC allele. In addition, under SSI, the results may vary depending on the dominance relationships between S-alleles (Novikova et al., 2023; Duan et al., 2024) and the level of inbreeding depression. As a result, additional factors and parameters would need to be examined compared to the current model.

## Acknowledgments

We thank Sylvain Billiard for helpful comments regarding the mathematical resolution of the neutral model. We also thank Boris Igic and Rosana Zenil-Ferguson for comments about empirical knowledge in the Solanaceae family. We thank the associate editor and two anonymous reviewers for their constructive comments.

## Author contributions

J.C. initiated the project. D.D. performed the mathematical analyses and developed the simulation model with guidance from J.C. and X.V. D.D. and J.C. wrote the first draft of the manuscript, and all authors edited it and approved the final version.

## Data availability

The simulation models are available at https://github.com/DianeDouet/GSI_autotetraploids

