## Supplementary Material for "Autopolyploid establishment under gametophytic self-incompatibility: the impact of self-fertilization and pollen limitation"

### A Supplementary Figures

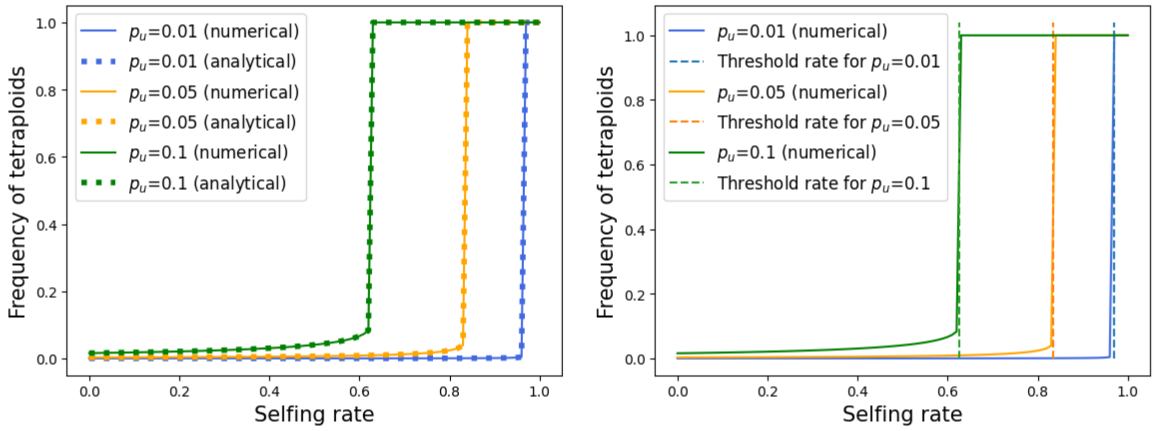


Figure S1: Frequency of tetraploids at equilibrium as a function of the selfing rate (left) for different values of $p_u$ obtained using the neutral model with a numerical resolution (full lines) and the analytical expressions from Equation 9 (dotted lines), and the threshold selfing rate at which tetraploids starts to establish into the population computed using Equation 8 (right).

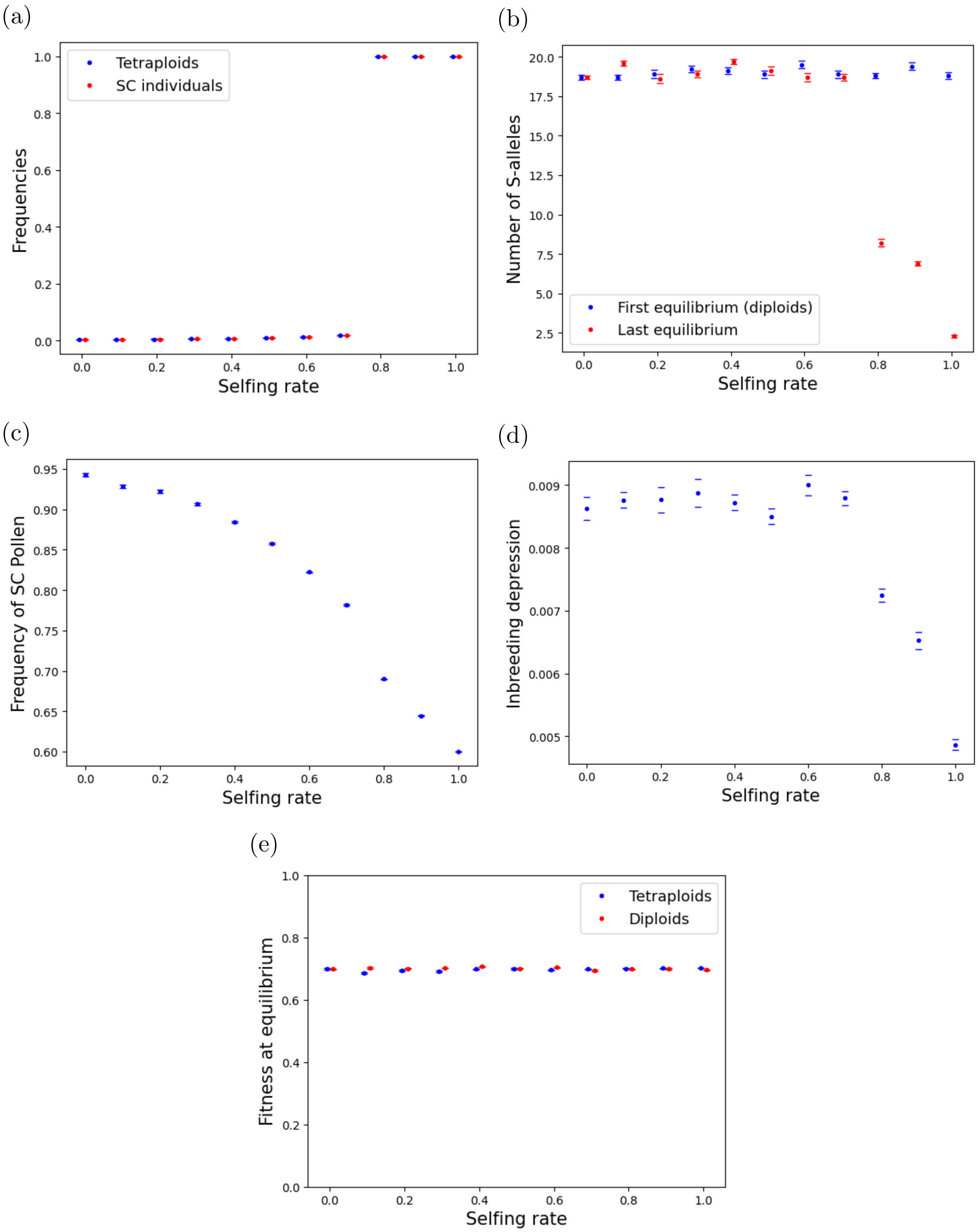


Figure S2: (a) Frequencies of tetraploids and SC individuals, (b) number of S-alleles at equilibrium for diploids and at the end of the simulation, (c) frequency of SC pollen produced by SC individuals, (d) inbreeding depression over the last 10000 generations, and (e) fitness of diploids and tetraploids at equilibrium when the selfing rate is fixed. Error bars stand for 95% confidence intervals. Results of 10 simulations runs under high pollen limitation (5 attempts) for each selfing rate using the parameters detailed in Table 1, specifically $N_{\text{pop}} = 1000$ and $p_u = 0.05$.

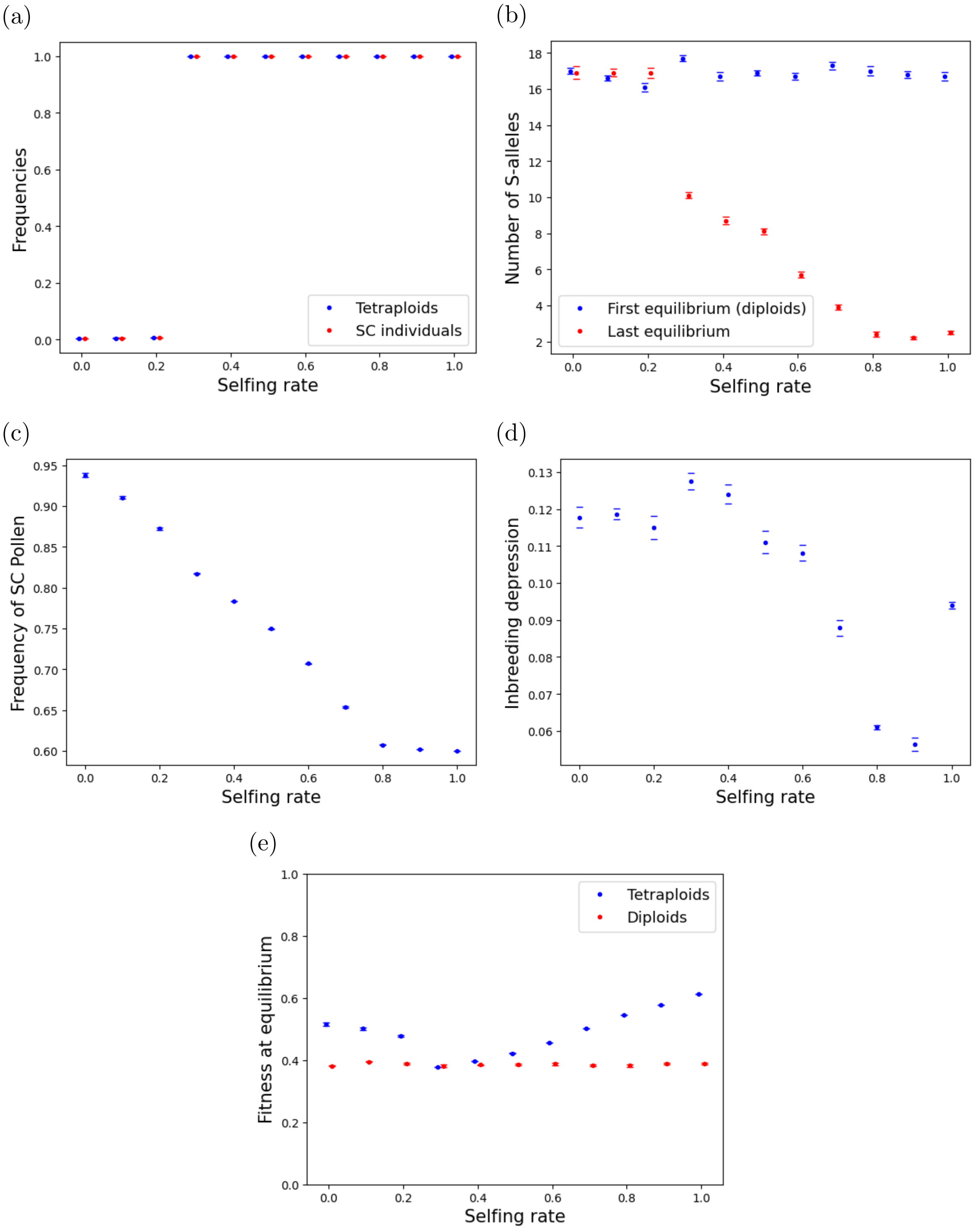


Figure S3: Evolutionary dynamics under high pollen limitation when the mutation rate is high ($U = 0.5$). (a) Frequencies of tetraploids and SC individuals, (b) number of S-alleles at equilibrium for diploids and at the end of the simulation, (c) frequency of SC pollen produced by SC individuals, (d) inbreeding depression over the last 10000 generations, and (e) fitness of diploids and tetraploids at equilibrium when the selfing rate is fixed. Error bars stand for 95% confidence intervals. Results of 10 simulations runs under high pollen limitation (1 attempts) for each selfing rate using the parameters detailed in Table 1, specifically $\omega^2 = 1$, $N_{\text{pop}} = 1000$ and $p_u = 0.05$.

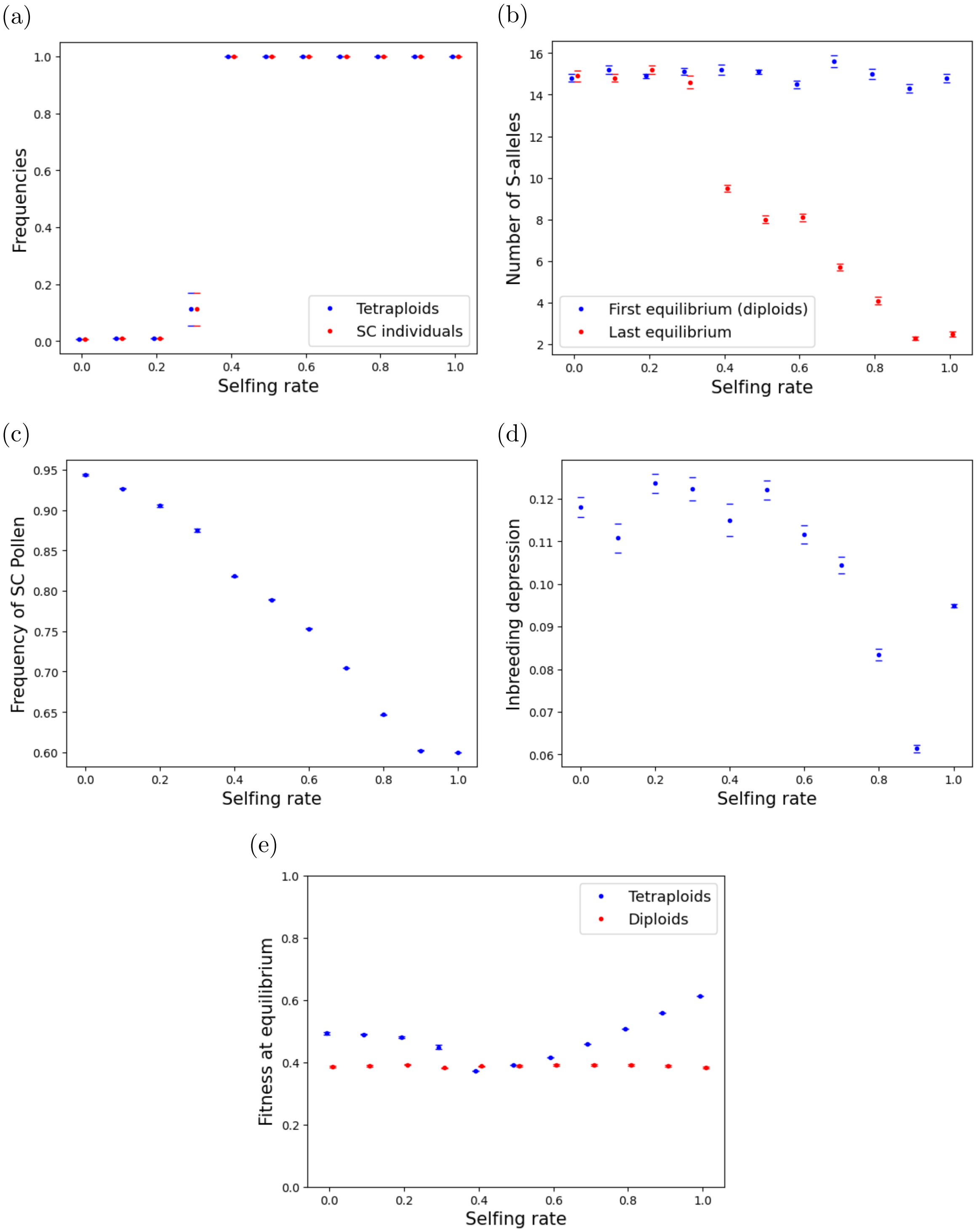


Figure S4: Evolutionary dynamics under low pollen limitation when the mutation rate is high ($U = 0.5$). (a) Frequencies of tetraploids and SC individuals, (b) number of S-alleles at equilibrium for diploids and at the end of the simulation, (c) frequency of SC pollen produced by SC individuals, (d) inbreeding depression over the last 10000 generations, and (e) fitness of diploids and tetraploids at equilibrium when the selfing rate is fixed. Error bars stand for 95% confidence intervals. Results of 10 simulations runs under low pollen limitation (20 attempts) for each selfing rate using the parameters detailed in Table 1, specifically $\omega^2 = 1$, $N_{\text{pop}} = 1000$ and $p_u = 0.05$.

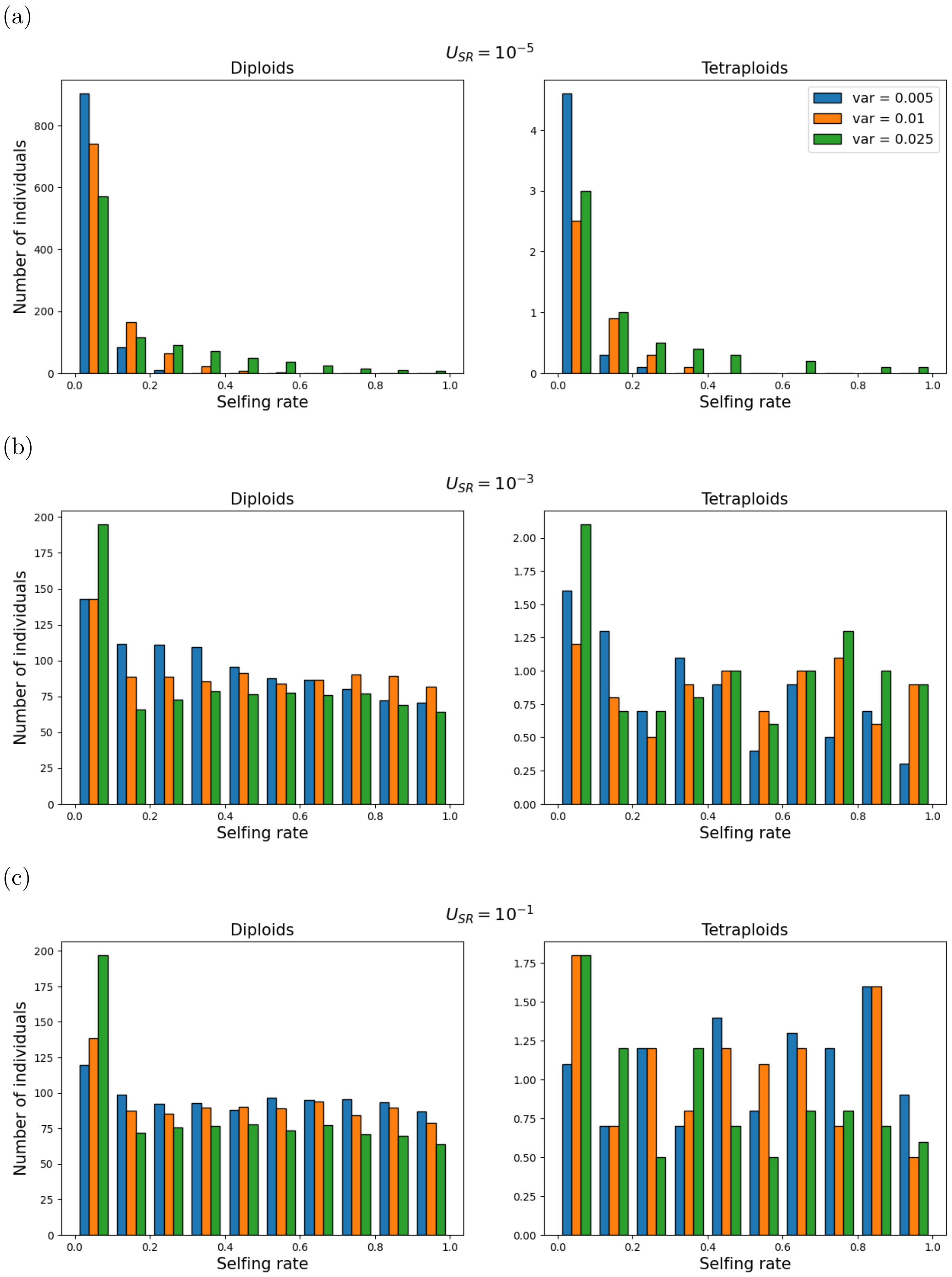


Figure S5: Distribution of the selfing rates of diploids (left) and tetraploids (right) at the end of the simulation for three different values of the variance $\sigma^2 = 0.05, 0.1$ and 0.25 and for different values of the mutation rate: (a) $U_{\mathrm{SR}} = 10^{-5}$, (b) $U_{\mathrm{SR}} = 10^{-3}$ and (c) $U_{\mathrm{SR}} = 10^{-1}$. Results averaged over 10 simulation runs under high pollen limitation (5 attempts) when the selfing rate can evolve using the parameters detailed in Table 1, specifically $N_{\mathrm{pop}} = 1000$ and $p_u = 0.05$.
